# Genome-wide characterization deciphers distinct properties of major intrinsic proteins in six *Phytophthora* species

**DOI:** 10.1101/394395

**Authors:** Abul Kalam Azad, Jahed Ahmed, Al Hakim, Md. Mahbub Hasan, Md. Asraful Alum, Mahmudul Hasan, Takahiro Ishikawa, Yoshihiro Sawa

## Abstract

Major intrinsic proteins (MIPs), commonly known as aquaporins, facilitate the membrane diffusion of water and some other non- polar solutes. MIPs might be involved in host-pathogen interactions. Herein, we identified 17, 24, 27, 19, 19, and 22 full-length MIPs, respectively, in the genomes of six *Phytophthora* species, *P. infestans, P. parasitica, P. sojae, P. ramorum, P. capsici*, and *P. cinnamomi*. These *Phytophthora* species are devastating plant pathogens and members of oomycetes, a distinct lineage of fungus-like eukaryotic microbes. Phylogenetic analysis showed that the *Phytophthora* MIPs (PMIPs) formed a completely distinct clade from their counterparts in other taxa and were clustered into nine subgroups. Sequence and structural properties indicated that the primary selectivity-related constrictions, including aromatic arginine (ar/R) selectivity filter and Froger’s positions in PMIPs were distinct from those in other taxa. The substitutions in the conserved Asn-Pro-Ala motifs in loops B and E of many PMIPs were also divergent from those in plants. We further deciphered group-specific consensus sequences/motifs in different loops and transmembrane helices of PMIPs, which were distinct from those in plants, animals, and microbes. The data collectively supported the notion that PMIPs might have novel functions.

## 1. Introduction

The superfamily of major intrinsic proteins (MIPs) possesses channel-forming integral membrane proteins that facilitate the transmembrane diffusion of water and other non-polar small solutes, such as ammonia, hydrogen peroxide, urea, boron, silicon, glycerol, carbon dioxide, antimony, and arsenite [1–8]. They are present in almost all living organisms from bacteria to mammals but highly abundant in plants [4, 5, 9]. Orthodox aquaporins (AQPs), which only facilitate the membrane diffusion of water and aquaglyceroporins (AQGPs), that can transport other uncharged small solutes in addition to water or without water, are prototype members of MIPs [3, 5, 10].

All AQPs share some structural features, however, their protein sequences differ substantially. Each monomer of MIP is constituted of six transmembrane (TM) α-helices (H1- H6) connecting with five loops (loops LA–LE) and forming an individual pore that theoretically specify the transport activity. There are two main constrictions in the channel. The first constriction is formed by two highly conserved Asn-Pro-Ala (NPA) motifs on loops B and E that is involved in proton exclusion [11, 12]. The second constriction, called the aromatic/arginine (ar/R) selectivity filter, is formed by four residues from helix 2 (H2), helix 5 (H5), and loop E (LE1 and LE2) [13, 14]. Mutation at this ar/R selectivity filter is thought to determine the broad spectrum of substrate conductance in plant AQPs [1, 2, 11, 15]. Additional five relatively conserved amino acid residues known as Froger’s positions and designed P1–P5, play roles in MIP sub-grouping and substrate transport selectivity [1, 16, 17].

Although 13 different MIPs identified in mammals are divided into three major subfamilies, the genomes of plants encode 2–5-fold or more MIPs, which are grouped into 4–7 subfamilies [1, 5, 18]. However, fungal genomes have up to five MIPs with diversified subgroups [19, 20]. Algal genomes have 1–6 MIPs, but are highly divergent and share only limited similarities with land plant MIPs [21]. In humans, MIPs play significant roles in brain-water balance, kidney nephrons, cell migration, cell proliferation, neural activity, pain, epithelial fluid secretion, skin hydration, adipocyte metabolism, and ocular function [6, 7]. They are associated with many human diseases, such as glaucoma, cancer, epilepsy, nephrogenic diabetes insipidus, and obesity [6, 7], and therefore, they have been potential drug targets [22]. In plants, MIPs have several vital physiological roles, from plant growth and development to physiological responses under biotic and abiotic stress conditions [4, 5, 23–31]. MIPs also play important roles in host-parasite interactions and are considered potential drug targets [32–37]. Recently, transcriptome data provided important clues about the participation of MIPs in host pathogen interactions [38–40]. In mycorrhizal plants, both plant and fungal MIPs play significant roles in nutrient uptake including water that actually helps to develop drought tolerant plants [5, 41, 42]. MIPs in pathogenic fungi may act as potential targets for antifungal drugs [20]. However, a systematic study of MIPs in oomycetes has not been done. Oomycetes are classified in the kingdom of Stramenopiles and include many economically important pathogens of plants, fish and aquatic invertebrates [43, 44].

Among oomycetes, the *Phytophthora* genus comprises more than 117 species, which are highly devastating to different kinds of agriculturally and ornamentally important plants, cause severe economic losses [45–50]. The present study includes only six *Phytophthora* species those have direct impact on agriculture by causing devastating plant diseases. The genome sequences of these six plant pathogenic *Phytophthora* species have been completed [45, 49, 51, 52] and are available in ‘FungiDB’ (http://fungidb.org/fungidb/). In this study, we identified and characterized *MIP* genes in the genomes of these six *Phytophthora* species (*PinMIP, PpaMIP, PsoMIP, PraMIP, PcaMIP*, and *PciMIP* genes of *P. infestans*, *P. parasitica*, *P. sojae, P. ramorum, P. capsici*, and *P. cinnamomi*, respectively) by using bioinformatics tools. To the best of our knowledge, this is the first report on MIPs of any organism of oomycetes. This report showed that the genomes of *Phytophthora* species have several-fold greater number of MIP homologues than those of algae, fungi, other parasites, and even more than in mammals. The numbers of their MIP homologues are comparable to that of plants. Comprehensive analysis with different bioinformatics tools revealed that the MIPs in *Phytophthora* species are phylogenetically and structurally distinct from their counterparts in other taxa.

## 2. Materials and methods

### 2.1. Identification of PinMIP, PpaMIP, PsoMIP, PraMIP, PcaMIP, and PciMIP genes

The genomes of *P. infestans* T30-4, *P. parasitica* INRA-310, *P. sojae* strain P6497*, P. ramorum* strain Pr102*, P. capsici* LT1534, and *P. cinnamomi* CBS 144.22 are available at FungiDB (http://fungidb.org/fungidb/). The AQPs protein sequences were retrieved from the FungiDB using a keyword “aquaporin” at gene text search option for above mentioned six *Phytophthora* species and checked the typical features of AQPs in their primary protein structures. All of the AQP protein sequences available in every *Phytophthora* species were used as queries for MIPs in the genome of a particular *Phytophthora* species using BLASTp tool with default parameters. Furthermore, the genomes of the *Phytophthora* species were searched for MIPs using BLASTp tool with the protein sequences of the complete set of 55 MIPs from *Populus trichocarpa* (PtMIP), 23 MIPs from *Physcomitrella patens* (PpMIP), and 12 MIPs from humans (MAQPs) as queries so that XIPs (uncharacterized X intrinsic proteins), GIPs (GlpF-like intrinsic proteins), HIPs (Hybrid intrinsic proteins) or superaquaporins could be identified if they were present in the genomes of the six *Phytophthora* species. MIPs in *P. infestans* (PinMIPs), *P. parasitica* (PpaMIPs), *P. sojae* (PsoMIPs)*, P. ramorum* (PraMIPs)*, P. capsici* (PcaMIPs), and *P. cinnamomi* (PciMIPs) were included until no more MIPs could be found from the corresponding species. All the sequences from each *Phytophthora* species were compared to identify the maximum number of MIPs for further analyses. Some of the MIP sequences might have been partial or might not have had all the features associated with its MIP channel. To investigate this, the multiple alignment program Clustal Omega (http://www.ebi.ac.uk/Tools/msa/clustalo/) was used to align all the sequences in an individual species. The multiple sequence alignment was used to determine the following features specific to the MIP family: (i) presence of two NPA or NPA-like motifs, (ii) presence of six TM α-helices, and (iii) two functionally important loops possessing the features characteristically present in MIP channels. The TM α-helices were predicted from the protein sequences of PMIPs by using SOSUI (http://bp.nuap.nagoya-u.ac.jp/sosui/), and TMpred (http://www.ch.embnet.org/software/TMPRED_form.html) servers. To determine the structure of *PMIP* genes, the gene and its CDS was used as input in the GeneMark.hmm ES-3.0 program [53], a self-training based algorithm for prediction of gene structures from novel eukaryotic genomes.

### 2.2. Phylogenetic analysis of PinMIPs, PpaMIPs, PsoMIPs, PraMIPs, PcaMIPs, and PciMIPs

Phylogenetic analysis was performed with MIPs in plants, animals, fungi, algae, bacteria, archaea and six *Phytophthora* species using Molecular Evolution Genetic Analysis (MEGA) version 5.0 [54] to understand the diversity and evolution of PMIPs. PinMIPs, PpaMIPs, PsoMIPs, PraMIPs, PcaMIPs, and PciMIPs were aligned with all PpMIPs and MAQPs, and 25 PtMIPs (5 homologues from each PtMIP subfamily), representative members from each of 10 subfamilies (90 homologues including 5-10 homologues from each subfamily) of MIPs in fungi [20], all MIPs in the genomes of nine algae [21], MIPs from five Gram-positive and seven Gram-negative bacteria and three archaea [55] using the Clustal Omega program (http://www.ebi.ac.uk/Tools/msa/clustalo/). A phylogenetic tree was generated with MEGA 5.0 using the maximum likelihood method. Reliability of individual branches of the tree was confirmed by performing bootstrapping with 1000 replicates. Another phylogenetic tree was constructed with all the MIPs in the six *Phytophthora* species using same parameters aforementioned. The identified PinMIPs, PpaMIPs, PsoMIPs, PraMIPs, PcaMIPs, and PciMIPs were divided into different subfamilies and groups based on the phylogenetic tree.

### 2.3. Prediction of subcellular localization and computation of Ka/Ks value for screening positive selection

The subcellular localizations of PMIPs were predicted *in silico* by using WoLF PSORT (http://www.genscript.com/wolf-psort.html) and Cello prediction system (http://cello.life.nctu.edu.tw/) as described previously [1].

To detect variation of selective pressure among PMIPs lineages, CODEML was used implemented in PAML package [56]. To determine whether the Ka/Ks ratio is significantly different from 1, a hypothesis, Ka/Ks=1 was tested using likelihood ratio test. To detect significant selective pressure among lineages, a null model (H_0_), Model 0 (one ratio model) assumes same ω (Ka/Ks) ratio for all lineages was compared to a parameter rich free-ratio model (H_A_, Model=1), where it assumes independent ω ratio for each branch.

### 2.4. Homology modeling

Homology models were generated using the Molecular Operating Environment software (MOE 2009.10; Chemical Computing Group, Quebec, Canada) as described previously [2]. The sequence of each MIP homolog was aligned with SoPIP2;1 (PDB2B5F) [57], or other AQP templates as indicated, using the MOE software as previously described [2]. The homology models were generated by using the MOE homology program and the stereochemical quality of the templates and models were assessed as previously described [9].

### 2.5. Determination of pore diameter and pore lining residues

The PoreWalker server (http://www.ebi.ac.uk/thornton-srv/software/PoreWalker/) [58], which is a fully automated method to analyze MIP channels, was used to detect and characterize PMIP channels from their 3D structures. The pore lining residues, which are involved in channel formation, were identified by using the PoreWalker server. The diameter of the narrowest part of the channel called the ar/R selectivity filter was also determined from the PoreWalker outputs as previously described [1].

## 3. Results

### 3.1. Genome-wide identification of PinMIP, PpaMIP, PsoMIP, PraMIP, PcaMIP, and PciMIP genes

The whole genome shotgun sequence (WGS) of *P. infestans*, *P. parasitica*, *P. sojae, P. ramorum, P. capsici*, and *P. cinnamomi* available at FungiDB was searched for *PinMIP, PpaMIP, PsoMIP, PraMIP, PcaMIP*, and *PciMIP* genes using BLASTp with default parameters. The query PinMIP, PpaMIP, PsoMIP, PtMIPs, PpMIPs and MAQPs sequences resulted in 21, 27, and 33 hits for *PinMIP, PpaMIP*, and *PsoMIP*, respectively. All the identified protein sequences of *PinMIP*, *PpaMIP*, and *PsoMIP* genes were manually inspected based on their TM domains and homology models. After these manual inspections, it was found that out of 21, 27, and 33 hits for *PinMIP, PpaMIP*, and *PsoMIP*, respectively, 4, 3, and 6 were considered to be pseudo *MIP* genes in *P. infestans*, *P. parasitica*, *P. sojae*, respectively, and were discarded. Characteristics, such as short sequences, N- or C-termini less, addition or deletion of sequences, and combinations thereof were observed in the discarded *MIPs*. However, the query PraMIP, PcaMIP, and PciMIP returned no hits in the BLASTp searches. Therefore, we retrieved the MIP sequences of *P. ramorum, P. capsici*, and *P. cinnamomi* from the FungiDB and analyzed them as mentioned above. Out of 32 MIP sequences in each of *P. ramorum, P. capsici*, and *P. cinnamomi*, 13 MIPs in the former two and 10 MIPs in the latter were deemed pseudo *MIPs* for the reasons noted above and were discarded. We ultimately obtained 17, 24, 27, 19, 19, and 22 full-length PinMIP, PpaMIP, PsoMIP, PraMIP, PcaMIP, and PciMIP protein sequences from the WGS of *P. infestans*, *P. parasitica*, *P. sojae, P. ramorum, P. capsici*, and *P. cinnamomi*, respectively (Tables 1–6). The amino acid lengths of PinMIP, PpaMIP, PsoMIP, PraMIP, PcaMIP, and PciMIP homologues with their maximum sequence identity with MIPs in other *Phytophthora*, humans (taxid 9606), *Arabidopsis thaliana* (taxid 3702), fungi (taxid 4751), and algae (taxid 3027) are tabulated in Tables 1–6. Although the MIPs of one *Phytophthora* species exhibited a maximum 80–95% identity with those of other *Phytophthora* species (Tables 1–6), the BLASTp search revealed that their highest identity with MIPs of *Arabidopsis thaliana* (taxid: 3702), *Homo sapiens* (taxid: 9606), fungi (taxid: 4751), and algae (taxid: 3027) was within 33, 45, 52, and 32%, respectively (Tables 1–6). This result indicated that PMIPs have higher identity with MIPs of fungi compared to those of animals, plants, and algae. This result further supported the notion that PMIPs might have extensive divergence in their sequence and structural properties compared to those in other taxonomic groups.

**Table 1.**
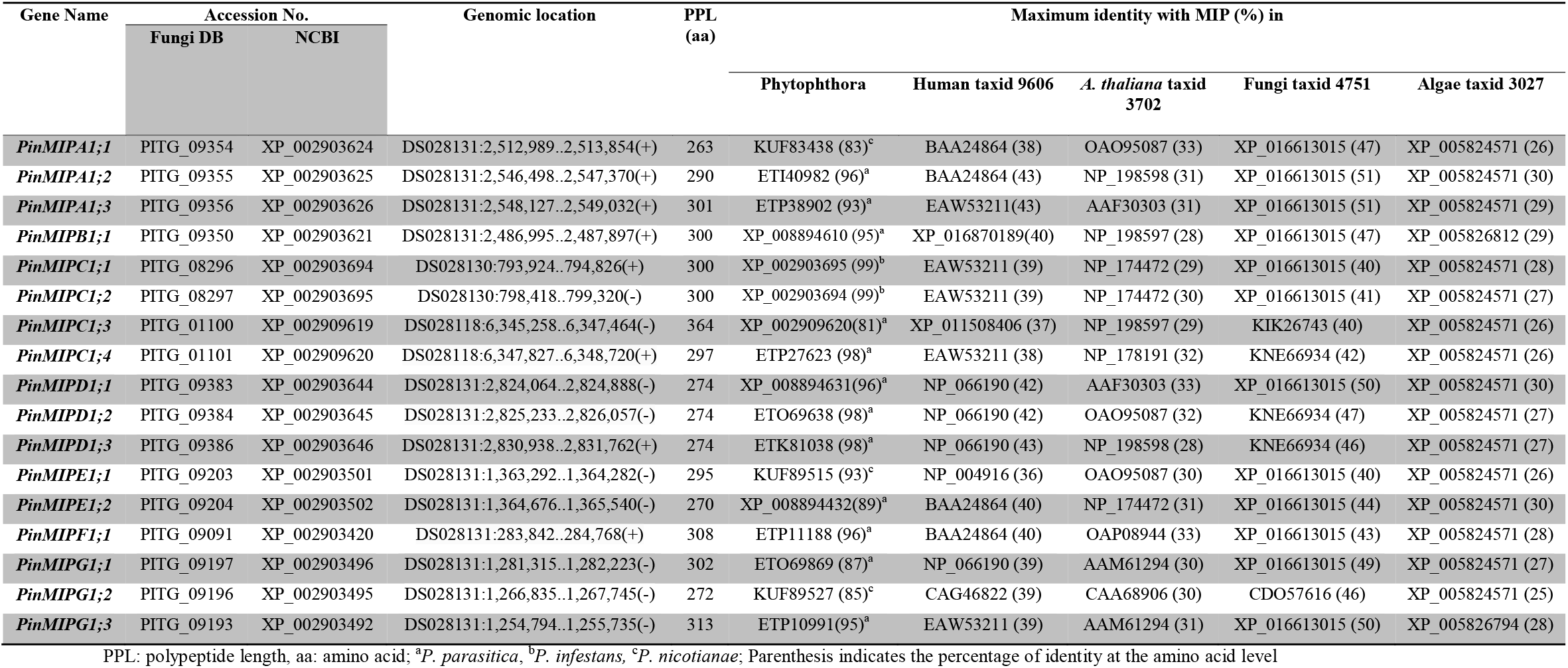
*MIP* genes in *P infestans* identified from the whole genome shotgun (WGS).

**Table 2.**
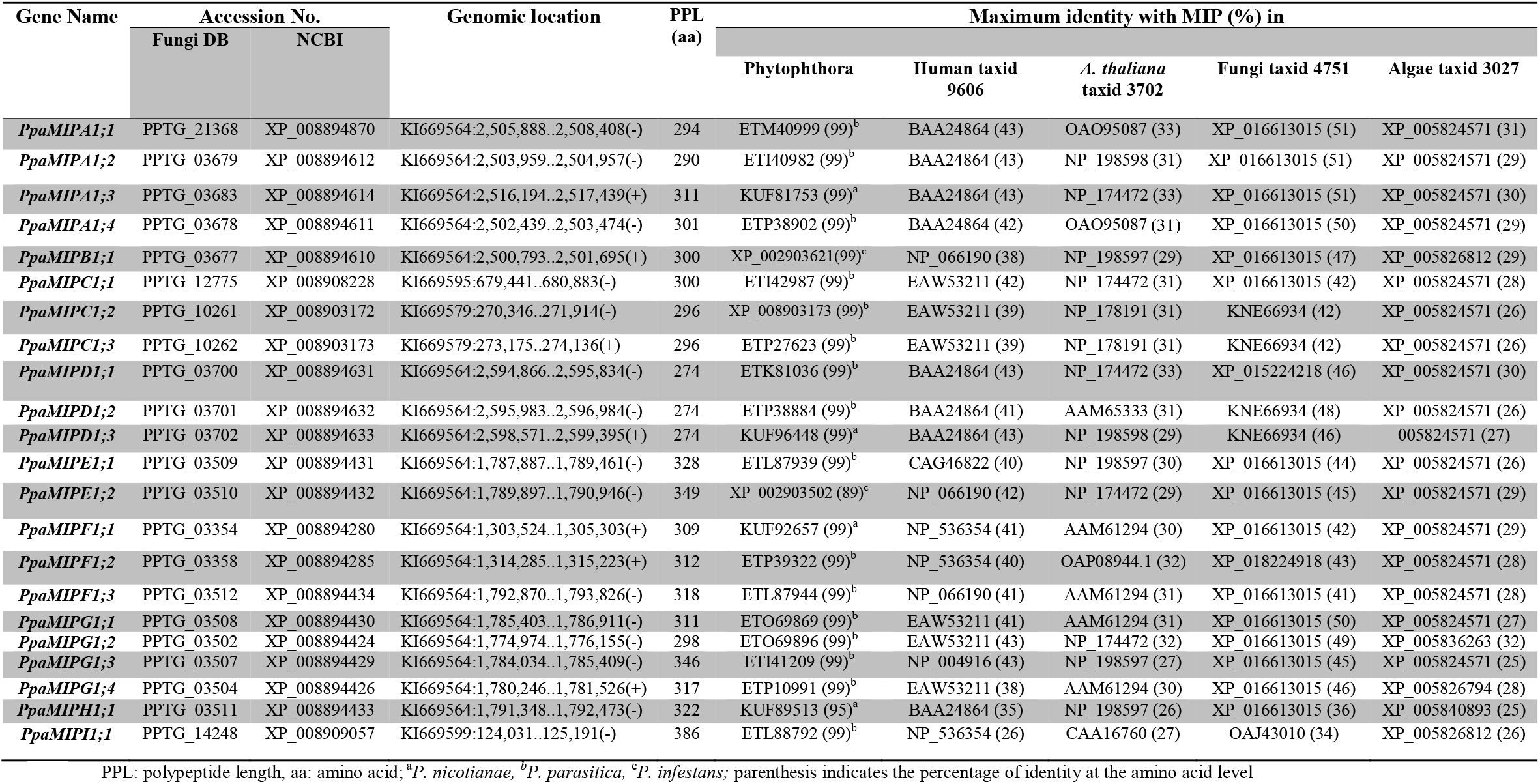
*MIP* genes in *P. parasitica* identified from the whole genome shotgun (WGS).

**Table 3.**
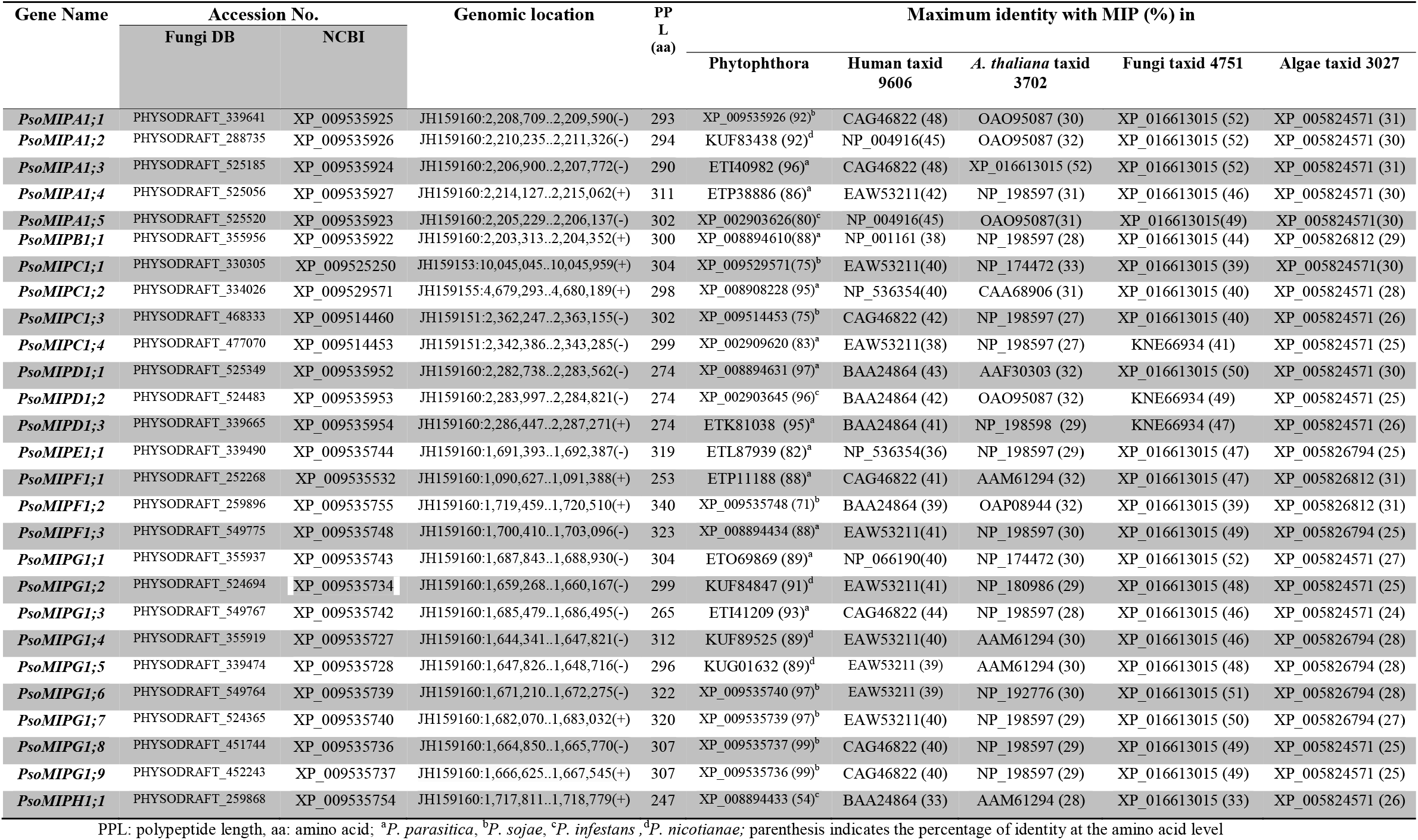
*MIP* genes in *P. sojae* identified from the whole genome shotgun (WGS).

| Gene Name | Accession No. |  | Genomic location | PP<br>L<br>(aa) | Maximum identity with MIP (%) in |  |  |  |  |
| --- | --- | --- | --- | --- | --- | --- | --- | --- | --- |
|  | Fungi DB | NCBI |  |  | Phytophthora | Human taxid<br>9606 | *A. thaliana* taxid<br>3702 | Fungi taxid 4751 | Algae taxid 3027 |
| *PsoMIPAI;1* | PHYSODRAFT_339641 | XP_009535925 | JH159160:2,208,709..2,209,590(-) | 293 | XP_009535926 (92)<sup>b</sup> | CAG46822 (48) | OAO95087 (30) | XP_016613015 (52) | XP_005824571 (31) |
| *PsoMIPAI;2* | PHYSODRAFT_288735 | XP_009535926 | JH159160:2,210,235..2,211,326(-) | 294 | KUF83438 (92)<sup>d</sup> | NP_004916(45) | OAO95087 (32) | XP_016613015 (52) | XP_005824571 (30) |
| *PsoMIPAI;3* | PHYSODRAFT_525185 | XP_009535924 | JH159160:2,206,900..2,207,772(-) | 290 | ETI40982 (96)<sup>a</sup> | CAG46822 (48) | XP_016613015 (52) | XP_016613015 (52) | XP_005824571 (31) |
| *PsoMIPAI;4* | PHYSODRAFT_525056 | XP_009535927 | JH159160:2,214,127..2,215,062(+) | 311 | ETP38886 (86)<sup>a</sup> | EAW53211(42) | NP_198597 (31) | XP_016613015 (46) | XP_005824571 (30) |
| *PsoMIPAI;5* | PHYSODRAFT_525520 | XP_009535923 | JH159160:2,205,229..2,206,137(-) | 302 | XP_002903626(80)<sup>c</sup> | NP_004916(45) | OAO95087(31) | XP_016613015(49) | XP_005824571(30) |
| *PsoMIPBI;1* | PHYSODRAFT_355956 | XP_009535922 | JH159160:2,203,313..2,204,352(+) | 300 | XP_008894610(88)<sup>a</sup> | NP_001161 (38) | NP_198597 (28) | XP_016613015 (44) | XP_005826812 (29) |
| *PsoMIPCI;1* | PHYSODRAFT_330305 | XP_009525250 | JH159153:10,045,045..10,045,959(+) | 304 | XP_009529571(75)<sup>b</sup> | EAW53211(40) | NP_174472 (33) | XP_016613015 (39) | XP_005824571(30) |
| *PsoMIPCI;2* | PHYSODRAFT_334026 | XP_009529571 | JH159155:4,679,293..4,680,189(+) | 298 | XP_008908228 (95)<sup>a</sup> | NP_536354(40) | CAA68906 (31) | XP_016613015 (40) | XP_005824571 (28) |
| *PsoMIPCI;3* | PHYSODRAFT_468333 | XP_009514460 | JH159151:2,362,247..2,363,155(-) | 302 | XP_009514453 (75)<sup>b</sup> | CAG46822 (42) | NP_198597 (27) | XP_016613015 (40) | XP_005824571 (26) |
| *PsoMIPCI;4* | PHYSODRAFT_477070 | XP_009514453 | JH159151:2,342,386..2,343,285(-) | 299 | XP_002909620 (83)<sup>a</sup> | EAW53211(38) | NP_198597 (27) | KNE66934 (41) | XP_005824571 (25) |
| *PsoMIPDI;1* | PHYSODRAFT_525349 | XP_009535952 | JH159160:2,282,738..2,283,562(-) | 274 | XP_008894631 (97)<sup>a</sup> | BAA24864 (43) | AAF30303 (32) | XP_016613015 (50) | XP_005824571 (30) |
| *PsoMIPDI;2* | PHYSODRAFT_524483 | XP_009535953 | JH159160:2,283,997..2,284,821(-) | 274 | XP_002903645 (96)<sup>c</sup> | BAA24864 (42) | OAO95087 (32) | KNE66934 (49) | XP_005824571 (25) |
| *PsoMIPDI;3* | PHYSODRAFT_339665 | XP_009535954 | JH159160:2,286,447..2,287,271(+) | 274 | ETK81038 (95)<sup>a</sup> | BAA24864 (41) | NP_198598 (29) | KNE66934 (47) | XP_005824571 (26) |
| *PsoMIPEI;1* | PHYSODRAFT_339490 | XP_009535744 | JH159160:1,691,393..1,692,387(-) | 319 | ETL87939 (82)<sup>a</sup> | NP_536354(36) | NP_198597 (29) | XP_016613015 (47) | XP_005826794 (25) |
| *PsoMIPFI;1* | PHYSODRAFT_252268 | XP_009535532 | JH159160:1,090,627..1,091,388(+) | 253 | ETP11188 (88)<sup>a</sup> | CAG46822 (41) | AAM61294 (32) | XP_016613015 (47) | XP_005826812 (31) |
| *PsoMIPFI;2* | PHYSODRAFT_259896 | XP_009535755 | JH159160:1,719,459..1,720,510(+) | 340 | XP_009535748 (71)<sup>b</sup> | BAA24864 (39) | OAP08944 (32) | XP_016613015 (39) | XP_005826812 (31) |
| *PsoMIPFI;3* | PHYSODRAFT_549775 | XP_009535748 | JH159160:1,700,410..1,703,096(-) | 323 | XP_008894434 (88)<sup>a</sup> | EAW53211(41) | NP_198597 (30) | XP_016613015 (49) | XP_005826794 (25) |
| *PsoMIPGI;1* | PHYSODRAFT_355937 | XP_009535743 | JH159160:1,687,843..1,688,930(-) | 304 | ETO69869 (89)<sup>a</sup> | NP_066190(40) | NP_174472 (30) | XP_016613015 (52) | XP_005824571 (27) |
| *PsoMIPGI;2* | PHYSODRAFT_524694 | XP_009535734 | JH159160:1,659,268..1,660,167(-) | 299 | KUF84847 (91)<sup>d</sup> | EAW53211(41) | NP_180986 (29) | XP_016613015 (48) | XP_005824571 (25) |
| *PsoMIPGI;3* | PHYSODRAFT_549767 | XP_009535742 | JH159160:1,685,479..1,686,495(-) | 265 | ETI41209 (93)<sup>a</sup> | CAG46822 (44) | NP_198597 (28) | XP_016613015 (46) | XP_005824571 (24) |
| *PsoMIPGI;4* | PHYSODRAFT_355919 | XP_009535727 | JH159160:1,644,341..1,647,821(-) | 312 | KUF89525 (89)<sup>d</sup> | EAW53211(40) | AAM61294 (30) | XP_016613015 (46) | XP_005826794 (28) |
| *PsoMIPGI;5* | PHYSODRAFT_339474 | XP_009535728 | JH159160:1,647,826..1,648,716(-) | 296 | KUG01632 (89)<sup>d</sup> | EAW53211 (39) | AAM61294 (30) | XP_016613015 (48) | XP_005826794 (28) |
| *PsoMIPGI;6* | PHYSODRAFT_549764 | XP_009535739 | JH159160:1,671,210..1,672,275(-) | 322 | XP_009535740 (97)<sup>b</sup> | EAW53211 (39) | NP_192776 (30) | XP_016613015 (51) | XP_005826794 (28) |
| *PsoMIPGI;7* | PHYSODRAFT_524365 | XP_009535740 | JH159160:1,682,070..1,683,032(+) | 320 | XP_009535739 (97)<sup>b</sup> | EAW53211(40) | NP_198597 (29) | XP_016613015 (50) | XP_005826794 (27) |
| *PsoMIPGI;8* | PHYSODRAFT_451744 | XP_009535736 | JH159160:1,664,850..1,665,770(-) | 307 | XP_009535737 (99)<sup>b</sup> | CAG46822 (40) | NP_198597 (29) | XP_016613015 (49) | XP_005824571 (25) |
| *PsoMIPGI;9* | PHYSODRAFT_452243 | XP_009535737 | JH159160:1,666,625..1,667,545(+) | 307 | XP_009535736 (99)<sup>b</sup> | CAG46822 (40) | NP_198597 (29) | XP_016613015 (49) | XP_005824571 (25) |
| *PsoMIPH1;1* | PHYSODRAFT_259868 | XP_009535754 | JH159160:1,717,811..1,718,779(+) | 247 | XP_008894433 (54)<sup>c</sup> | BAA24864 (33) | AAM61294 (28) | XP_016613015 (33) | XP_005824571 (26) |

PPL: polypeptide length, aa: amino acid; <sup>a</sup>*P. parasitica*, <sup>b</sup>*P. sojae*, <sup>c</sup>*P. infestans*, <sup>d</sup>*P. nicotianae*; parenthesis indicates the percentage of identity at the amino acid level

**Table 4.** *MIP* genes in *P. ramorum* identified from the whole genome shotgun (WGS).

| Gene Name | Accession No. |  | Genomic location | PPL<br>(aa) | Maximum identity with MIP (%) in |  |  |  |  |
| --- | --- | --- | --- | --- | --- | --- | --- | --- | --- |
|  | Fungi DB | N<br>C<br>BI |  |  | Phytophthora | Human taxid<br>9606 | *A. thaliana* taxid<br>3702 | Fungi taxid 4751 | Algae taxid 3027 |
| *PraMIPAI;1* | PSURA_84475 | - | PramPr-102_SC0114:47,366..48,247(-) | 293 | KUF83438 (92)<sup>d</sup> | BAA24864 (41) | NP_198598 (31) | XP_016613015 (31) | XP_005824571 (29) |
| *PraMIPAI;2* | PSURA_84474 | - | PramPr-102_SC0114:45,578..46,447(-) | 289 | ETI40982 (96)<sup>a</sup> | BAA24864 (35) | NP_198598 (31) | XP_016613015 (49) | XP_005824571 (27) |
| *PraMIPAI;3* | PSURA_84476 | - | PramPr-102_SC0114:50,260..51,177(-) | 305 | ETP38886.1 (88)<sup>a</sup> | EAW53211 (41) | NP_198597 (30) | XP_016613015 (45) | XP_005826794 (27) |
| *PraMIPBI;1* | PSURA_72233 | - | PramPr-102_SC0114:42,371..43,273(+) | 300 | XP_008894610 (94)<sup>a</sup> | NP_066190 (38) | NP_198597 (29) | XP_016613015 (46) | XP_005826812 (30) |
| *PraMIPCI;1* | PSURA_71169 | - | PramPr-102_SC0006:392,090..392,875(+) | 261 | XP_008908228 (94)<sup>a</sup> | EAW53211 (40) | AAM61294 (31) | XP_016613015 (41) | XP_005824571 (27) |
| *PraMIPCI;2* | PSURA_72183 | - | PramPr-102_SC0102:39,914..40,669(-) | 251 | XP_009514460 (94)<sup>b</sup> | CAG46822 (41) | AAM61294 (31) | XP_016613015 (39) | XP_005824571 (25) |
| *PraMIPCI;3* | PSURA_72182 | - | PramPr-102_SC0102:33,841..34,596(+) | 251 | XP_009514453 (84)<sup>b</sup> | CAG46822 (41) | AAM61294 (30) | KNE66934 (42) | XP_005824571 (28) |
| *PraMIPDI;1* | PSURA_72235 | - | PramPr-102_SC0114:104,901..105,626(-) | 241 | XP_009535952 (96)<sup>b</sup> | BAA24864 (44) | NP_174472 (33) | XP_016613015 (50) | XP_005824571 (30) |
| *PraMIPDI;2* | PSURA_72236 | - | PramPr-102_SC0114:106,106..106,930(-) | 274 | XP_002903645 (96)<sup>c</sup> | NP_066190 (40) | OAO95087 (31) | XP_016613015 (49) | XP_005824571 (26) |
| *PraMIPDI;3* | PSURA_72237 | - | PramPr-102_SC0114:108,512..109,339(+) | 275 | XP_009535954 (95)<sup>b</sup> | NP_066190 (41) | NP_198598 (29) | KNE66934 (47) | XP_005824571 (26) |
| *PraMIPEI;1* | PSURA_80174 | - | PramPr-102_SC0046:362,803..363,804(+) | 333 | XP_008894431 (87)<sup>a</sup> | BAA24864 (40) | NP_198597 (32) | XP_016613015 (50) | XP_005824571 (26) |
| *PraMIPFI;1* | PSURA_95006 | - | PramPr-102_SC0029:82,759..83,706(+) | 315 | ETP11188 (80)<sup>a</sup> | BAA24864 (39) | AAM61294 (31) | XP_016613015 (47) | XP_005824571 (27) |
| *PraMIPFI;2* | PSURA_80164 | - | PramPr-102_SC0046:326,376..327,332(+) | 318 | ETP11188 (83)<sup>a</sup> | BAA24864 (38) | AAM61294 (31) | XP_016613015 (44) | XP_005824571 (28) |
| *PraMIPGI;1* | PSURA_71778 | - | PramPr-102_SC0046:367,948..368,772(-) | 274 | ETO69869 (93)<sup>a</sup> | NP_536354 (42) | AAM61294 (30) | XP_016613015 (51) | XP_005824571 (27) |
| *PraMIPGI;2* | PSURA_87677 | - | PramPr-102_SC2368:1,241..2,140(-) | 299 | XP_009535734 (93)<sup>b</sup> | EAW53211 (41) | NP_180986 (31) | XP_016613015 (49) | XP_005826794 (25) |
| *PraMIPGI;3* | PSURA_72426 | - | PramPr-102_SC0716:5,808..6,707(-) | 299 | XP_009535734 (93)<sup>b</sup> | EAW53211(41) | NP_180986 (31) | XP_016613015 (49) | XP_005824571 (25) |
| *PraMIPGI;4* | PSURA_71777 | - | PramPr-102_SC0046:365,813..366,649(-) | 278 | ETI41209 (91)<sup>a</sup> | CAG46822 (42) | NP_198597 (31) | XP_016613015 (46) | XP_005841067 (30) |
| *PraMIPGI;5* | PSURA_72419 | - | PramPr-102_SC0660:6,538..7,374(+) | 278 | KUF89525.1 (91)<sup>d</sup> | EAW53211(41) | AAM61294(29) | XP_016613015 (48) | XP_005826794 (28) |
| *PraMIPGI;6* | PSURA_72425 | - | PramPr-102_SC0716:955..1,704(-) | 249 | XP_009535728 (92)<sup>b</sup> | NP_004916 (43) | NP_192776 (31) | XP_016613015 (49) | XP_005826794 (28) |

PPL: polypeptide length, aa: amino acid; <sup>a</sup>*P. parasitica*, <sup>b</sup>*P. sojae*, <sup>c</sup>*P. infestans*, <sup>d</sup>*P. nicotianae*; Parenthesis indicates the percentage of identity at the amino acid level

**Table 5.** *MIP* genes in *P. capsici* identified from the whole genome shotgun (WGS).

| Gene Name | Accession No. |  | Genomic location | PPL<br>(aa) | Maximum identity with MIP (%) in |  |  |  |  |
| --- | --- | --- | --- | --- | --- | --- | --- | --- | --- |
|  | Fungi DB | N<br>C<br>B<br>I |  |  | Phytophthora | Human taxid<br>9606 | *A. thaliana* taxid<br>3702 | Fungi taxid 4751 | Algae taxid 3027 |
| *PcaMIPAI;1* | PHYCA_530062 | - | PcapLT1534_SC054:141,263..142,555(-) | 293 | KUF83438 (95)<sup>d</sup> | CAG46822 (45) | OAO95087 (32) | XP_016613015 (51) | XP_005824571 (32) |
| *PcaMIPAI;2* | PHYCA_510138 | - | PcapLT1534_SC054:139,634..140,630(-) | 288 | XP_009535925 (94)<sup>b</sup> | CAG46822 (44) | OAO95087 (33) | XP_016613015 (51) | XP_005824571 (31) |
| *PcaMIPAI;3* | PHYCA_553594 | - | PcapLT1534_SC054:138,017..138,944(-) | 291 | XP_009535924 (96)<sup>b</sup> | NP_066190 (43) | OAO95087 (32) | XP_016613015 (51) | XP_005824571 (31) |
| *PcaMIPAI;4* | PHYCA_553588 | - | PcapLT1534_SC054:136,640..137,545(-) | 301 | ETP38902 (90)<sup>a</sup> | EAW53211 (43) | NP_178191 (32) | XP_016613015 (48) | XP_005824571 (30) |
| *PcaMIPCI;1* | PHYCA_6911 | - | PcapLT1534_SC015:427,750..438,233(-) | 280 | XP_008908228 (87)<sup>a</sup> | EAW53211 (35) | NP_567572 (30) | XP_016613015 (39) | XP_005824571 (29) |
| *PcaMIPCI;2* | PHYCA_123018 | - | PcapLT1534_SC049:168,741..169,683(+) | 297 | XP_009514460 (90)<sup>b</sup> | EAW53211 (38) | NP_198597 (26) | XP_016613015 (40) | XP_005824571 (26) |
| *PcaMIPCI;3* | PHYCA_509760 | - | PcapLT1534_SC049:192,973..193,950(+) | 299 | XP_002909620 (91)<sup>c</sup> | EAW53211 (39) | NP_198597 (27) | KNE66934 (40) | XP_005824571 (26) |
| *PcaMIPDI;1* | PHYCA_553631 | - | PcapLT1534_SC054:213,802..214,626(-) | 274 | XP_009535952 (97)<sup>b</sup> | BAA24864 (42) | AAF30303 (32) | XP_016613015 (50) | XP_005824571 (30) |
| *PcaMIPDI;2* | PHYCA_510151 | - | PcapLT1534_SC054:214,814..215,848(-) | 274 | XP_002903645 (97)<sup>c</sup> | NP_066190 (42) | NP_198597 (33) | KNE66934 (48) | XP_005824571 (27) |
| *PcaMIPDI;3* | PHYCA_124758 | - | PcapLT1534_SC054:222,559..223,383(+) | 274 | ETK81038 (96)<sup>a</sup> | BAA24864 (42) | NP_001330234 (29) | KNE66934 (47) | XP_005824571 (27) |
| *PcaMIPEI;1* | PHYCA_130684 | - | PcapLT1534_SC096:30,774..31,649(-) | 291 | XP_008894431 (95)<sup>a</sup> | CAG46822 (41) | OAO95087(31) | XP_016613015 (46) | XP_005824571 (27) |
| *PcaMIPEI;2* | PHYCA_130620 | - | PcapLT1534_SC096:32,191..33,045(-) | 284 | XP_008894432 (92)<sup>a</sup> | BAA24864 (39) | NP_174472 (29) | XP_016613015 (46) | XP_005824571 (28) |
| *PcaMIPFI;1* | PHYCA_125677 | - | PcapLT1534_SC059:215,629..216,567(+) | 312 | ETP11193 (88)<sup>a</sup> | NP_001161 (38) | OAP08944 (33) | XP_016613015 (41) | XP_005824571 (30) |
| *PcaMIPFI;2* | PHYCA_130673 | - | PcapLT1534_SC096:35,031..35,984(-) | 317 | XP_008894434 (86)<sup>a</sup> | BAA24864 (42) | AAM61294 (29) | XP_016613015 (42) | XP_005826812 (31) |
| *PcaMIPGI;1* | PHYCA_511729 | - | PcapLT1534_SC096:27,663..28,733(-) | 274 | ETO69869 (93)<sup>a</sup> | EAW53211 (41) | AAM61294 (32) | XP_016613015 (52) | XP_005824571 (28) |
| *PcaMIPGI;2* | PHYCA_511737 | - | PcapLT1534_SC096:84,419..89,905(-) | 300 | XP_009535734 (92)<sup>b</sup> | EAW53211 (41) | AAM61294 (31) | XP_016613015 (51) | XP_005824571 (25) |
| *PcaMIPGI;3* | PHYCA_511725 | - | PcapLT1534_SC096:15,964..17,034(-) | 304 | ETI41209 (87)<sup>a</sup> | NP_004916 (39) | CAA68906 (29) | XP_016613015 (44) | XP_005841067 (32) |
| *PcaMIPGI;4* | PHYCA_512201 | - | PcapLT1534_SC669:271..1,219(+) | 297 | XP_009535 27 (80)<sup>b</sup> | EAW53211 (38) | AAM61294 (29) | CDO57616 (46) | XP_005826794 (28) |
| *PcaMIPGI;5* | PHYCA_130634 | - | PcapLT1534_SC096:96,478..97,393(+) | 268 | XP_009535727 (85)<sup>b</sup> | EAW53211 (38) | NP_192776 (27) | XP_016613015 (42) | XP_005826794 (27) |

PPL: polypeptide length, aa: amino acid; <sup>a</sup>*P. parasitica*, <sup>b</sup>*P. sojae*, <sup>c</sup>*P. infestans*, <sup>d</sup>*P. nicotianae*; parenthesis indicates the percentage of identity at the amino acid level

**Table 6.**
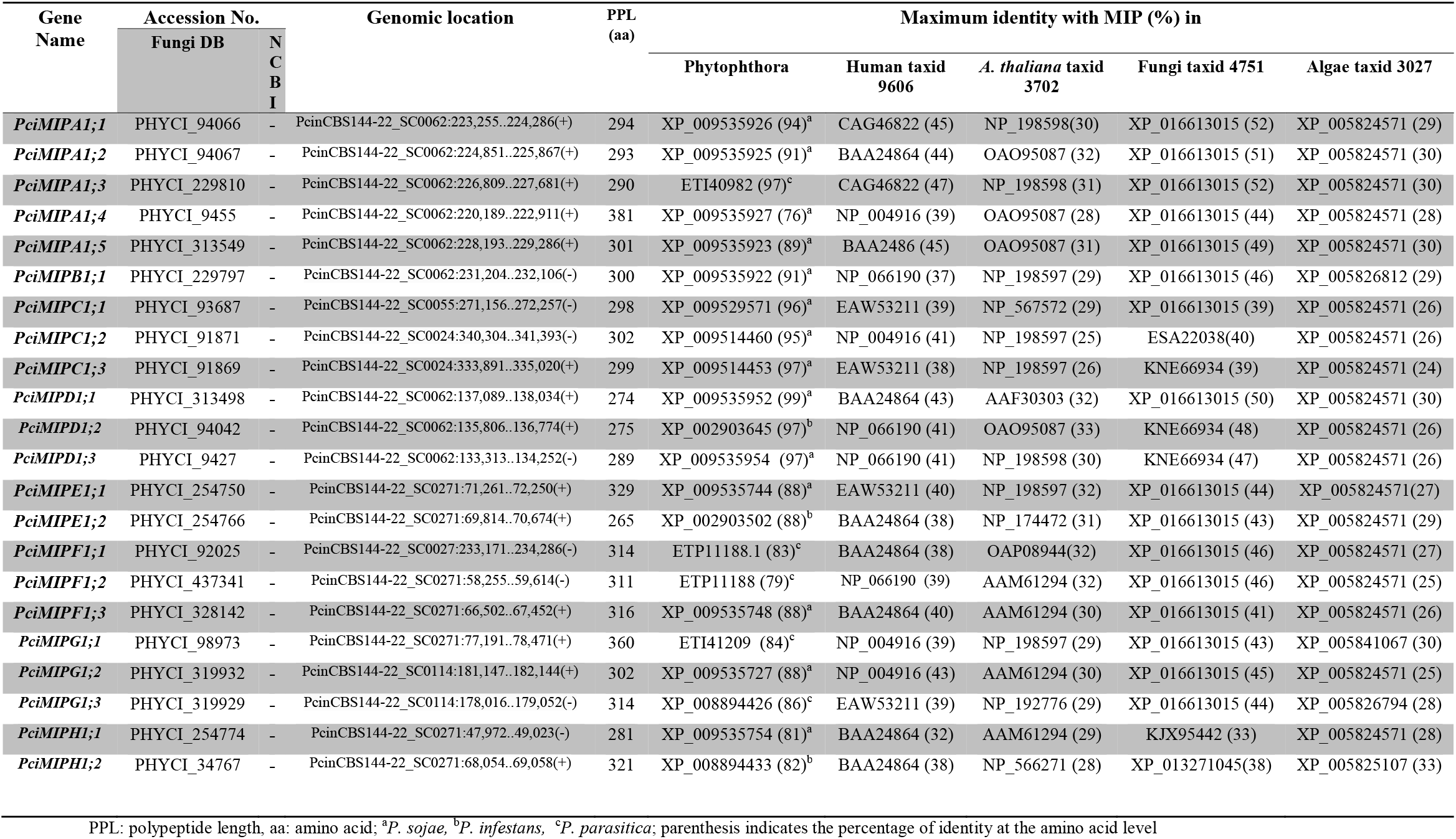
*MIP* genes in *P. cinnamomi* identified from the whole genome shotgun (WGS).

The score of the Cello prediction system showed that all PMIPs might have been localized in the plasma membrane (Table S1). However, WoLF PSORT scores suggested that some PMIPs have possibility to localize in other intragranular membranes, such as the endoplasmic reticulum, vacuole, golgi, and mitochondria.

Measuring the non-synonymous substitution (Ka) over synonymous substitution (Ks) rate is an important way to understand the molecular sequence evolution. The overabundance of non-synonymous substitutions rate over synonymous rate ratio (Ka/Ks or ω) is indicator of positive Darwinian selection. The ω value significantly greater than 1 means positive selection [59], where Ka is the number of nonsynonymous substitutions per nonsynonymous site and Ks is the number of synonymous substitutions per synonymous site [60, 61]. Thus, if an amino acid change is neutral, it will be fixed at the same rate as a synonymous mutation, with ω = 1. If the amino acid change is deleterious, purifying selection will reduce its fixation rate so that ω < 1. When the amino acid change offers a selective advantage, it will be fixed at a higher rate than a synonymous mutation, with ω > 1 [62].

To detect significant selective pressure among *PMIP* lineages, maximum likelihood approach followed one ratio model (null model, H_0_), was compared to an alternative (H_1_) free-ratio model [63]. The log likelihood values were calculated if H_A_ was significant over H_0_. One ratio model estimated log-likelihood value ℓ_0_ = −17290.60063, and ω = 0.54737. This ω ratio was an average over all lineages. The free-ration model was applied to the same data set. Log-likelihood value estimated by free-ratio model was ℓ_1_ = −17123.327460. Twice the difference between log-likelihood of two models (2Δℓ = 2(ℓ_1_ – ℓ_0_)) is 334.546342. The statistics with chi-square distribution (df=248) accept the free ratio model, with p-value 0.000201982. The free ratio model showed that 24 *PMIPs* in six *Phytophthora* species have ω >1, indicating that the evolution of these genes was likely under positive or Darwinian selection (Supplementary Fig. 1). The most of *PMIP* lineages showed ω values <1, indicating purifying selection. Pairwise comparison among PMIPs showed Ks > Ka, which were also indicative of purifying selection.

### 3.2. Phylogenetic distribution and nomenclature of PMIPs

The sequences of PMIPs were routinely compared with those in the MIP subfamilies of other kingdoms by constructing multiple and/or pairwise alignments using Clustal Omega and EMBOSS Needle, respectively. To investigate the structural conservation of PMIPs with other MIPs in plants, animals, and microbes, a 3D structural alignment was constructed with the templates of human AQP1 (PDB ID, 1J4N), spinach plasma membrane intrinsic protein, SoPIP2;1 (PDB ID, 2B5F), and *E. coli* glycerol facilitator, GlpF (PDB ID, 1FX8). All 3D models constructed with these three templates showed the typical hourglass shaped AQPs with six TM helices and five connecting loops (Fig. 1). Superposition of the three models of each PMIP showed that the helices superposed very closely. Deviation was observed only in the loops. This structural alignment was used as a guide for sequence alignment. To classify the PMIPs, their protein sequences were analyzed phylogenetically with MIPs in the plant, human, fungi, algae, bacteria and archaea, as described in Materials and Methods. Although PtMIPs and PpMIPs were divided into five and seven subfamilies, respectively [15, 64], and those in fungi and algae were separated into ten and seven subfamilies, respectively [20, 21], PMIPs formed a completely distinct clade (Fig. 2). Despite one MIP in *P. parasitica* (PpaMIPI1;1) which formed a distinct clade, no other PMIP was associated with any subfamilies of MIPs in other organisms in the three domains of life. Therefore, we generated a phylogenetic tree with all 126 MIP homologues in the six *Phytophthora* species (Fig. 3 and Supplementary Fig. 2). The 126 MIPs were clustered into nine subgroups. Because the PMIPs were not phylogenetically clustered with any subfamily or group of MIPs reported in plants, animals, fungi, algae, bacteria or archaea, these groups were arbitrarily named MIPA-MIPI. Each MIP homologue in every *Phytophthora* species was named by taking the first letter (Upper case) from the genus and the first two letters (lower case) of the species names with the MIP group (A to I), and the number of the homologues in each group was stated consecutively, i.e., PsoMIPA1;1 for MIPA in *P. sojae* and so on. MIPAs and MIPCs-MIPGs included one to several MIPs from each of the six *Phytophthora* species. MIPBs contained five homologues of five species, except *P. capsici*, and MIPHs consisted of four MIPs from three *Phytophthora* species. The MIPI group had only one homologue from *P. parasitica*, and merely this homologue clustered with delta aquaglyceroporins of fungi (Fig. 2). However, although PMIPB, PMIPH and PMIPI included a small number of MIP homologues, they had a distinct ar/R constriction. The characteristics of each of the nine PMIPs groups are detailed in the next sections.

**Fig. 1.**
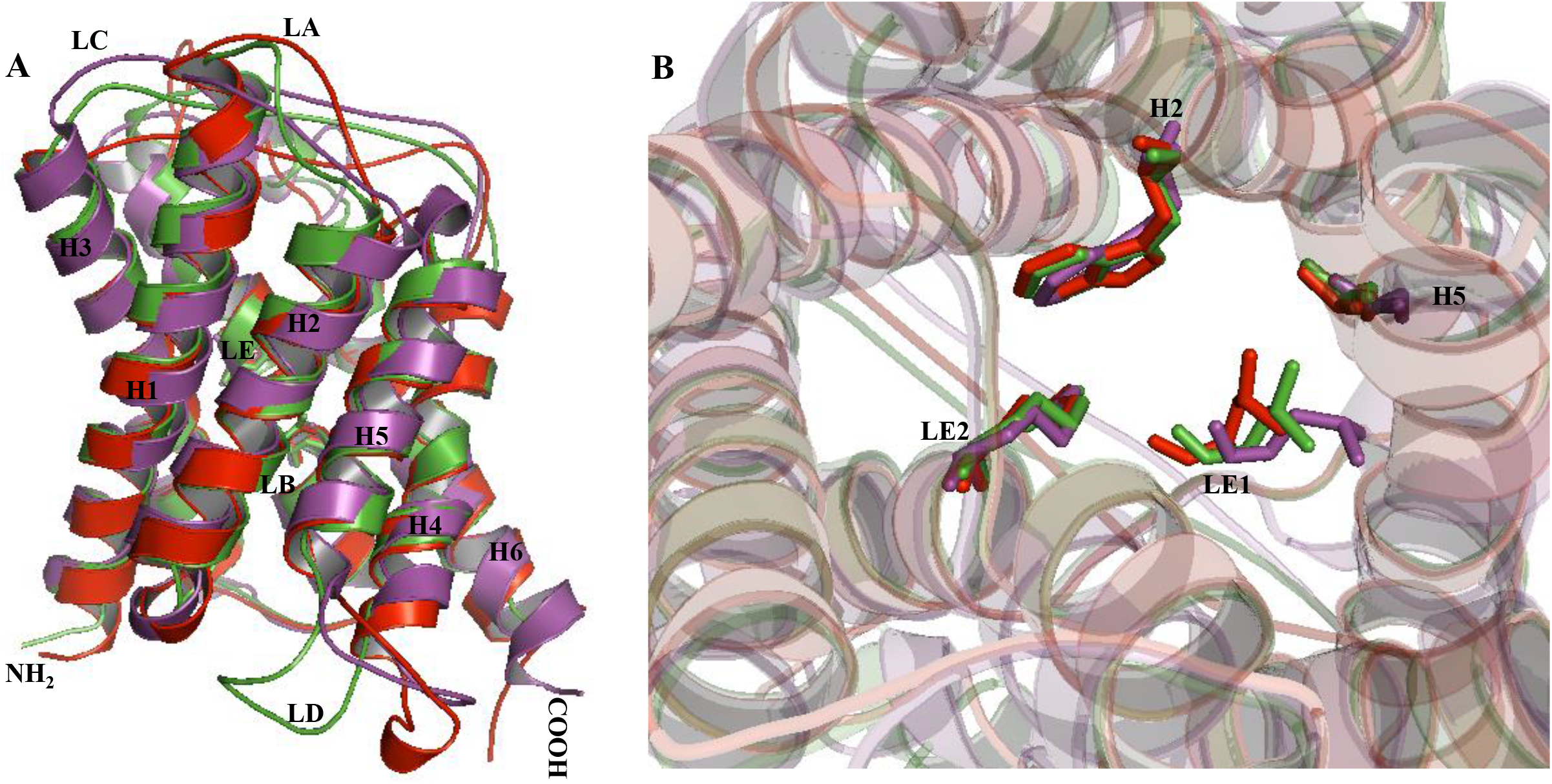
Superposition of representative 3D models of PcaMIPC1;1 constructed with the template AQP1 (red), GlpF (violet), and SoPIP2;1 (green). A and B are side and top views of the superposed models, respectively. The 3D models of all PMIPs were first constructed separately with AQP1, GlpF, and SoPIP2;1, and then the model of PcaMIPC1;1 as representative of PMIPs was superposed. The TM α-helices (H1-H6) and loops (LA-LE) are indicated. The NPA motifs in LB and LE are shown as sticks. The residues that form the aromatic/arginine (ar/R) tetrad in the superposed structures are shown as sticks (B). The TM α-helices and the loops to which these residues belong are indicated.

**Fig. 2.**
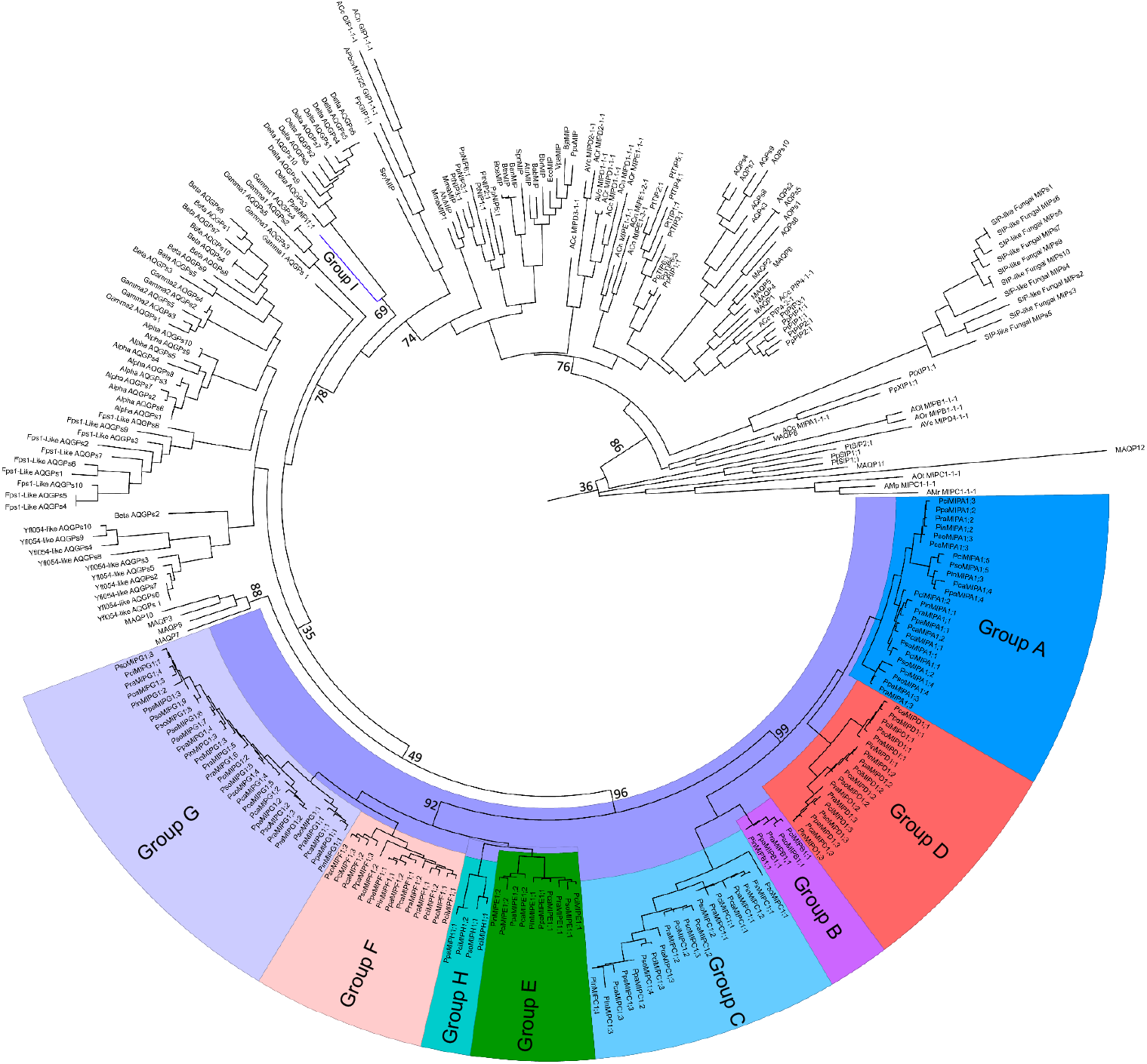
Evolutionary relationship of MIPs in the six *Phytophthora* species. Phylogenetic analysis of all PMIPs from the six *Phytophthora* species is shown along with MIPs from *Populus trichocarpa* (PtMIPs; PtPIPs, PtTIPs, PtNIPs, PtSIPs, and PtXIPs), *Physcomitrella patens* (PpMIPs), and human (MAQP), and representative members from each of 10 subfamilies of MIPs in fungi [20] (90 homologues including 5-10 homologues from each subfamily namely AQPs, Alpha AQGPs, Beta AQGPs, Gamma 1 AQGPs, Gamma 2 AQGPs, Delta AQGPs, Froger’s positions like AQGPs, XIPs, SIP-like Fungus MIP, and Yfl054-like AQGPs), all MIPs in the genomes of nine algae [21] (all homologues start with ‘A’), 12 bacterial MIPs (EcoMIP, *E. coli;* VpaMIP, *V. parahaemolyticus;* BbrMIP, *B. bronchiseptica;* AtuMIP, *A. tumefaciens;* BabMIP, *B. abortus;* BjaMIP, *B. japonicum;* PpuMIP, *P. pudida;* SpnMIP, *S. pneumonia;* BanMIP, *B. anthracis*; BceMIP, *B. cereus*; BthMIP, *B. thuringiensis*; SpyMIP, *S. pyogenes*) and 3 archaeal MIPs (AfuMIP, *A. fulgidus*; MmaMIP, *M. marburgensise*; MmaMIP1, *M. maripulodis)* [55]. The deduced amino acid sequences of MIPs were aligned using the Clustal Omega computer program and a phylogenetic tree was constructed using MEGA. The evolutionary history was inferred using the Bootstrap maximum likelihood (1000 replicates) method. PMIPs are shown with different background colors. All PMIPs, except PpaMIPI1;1, formed a distinct clade and did not cluster with any subfamily or group of MIPs in plants, humans, fungi, algae, bacteria or archaea. The PpaMIPI1;1 shown with the blue line clustered with the delta aquaglyceroporins of fungi.

**Fig. 3.**
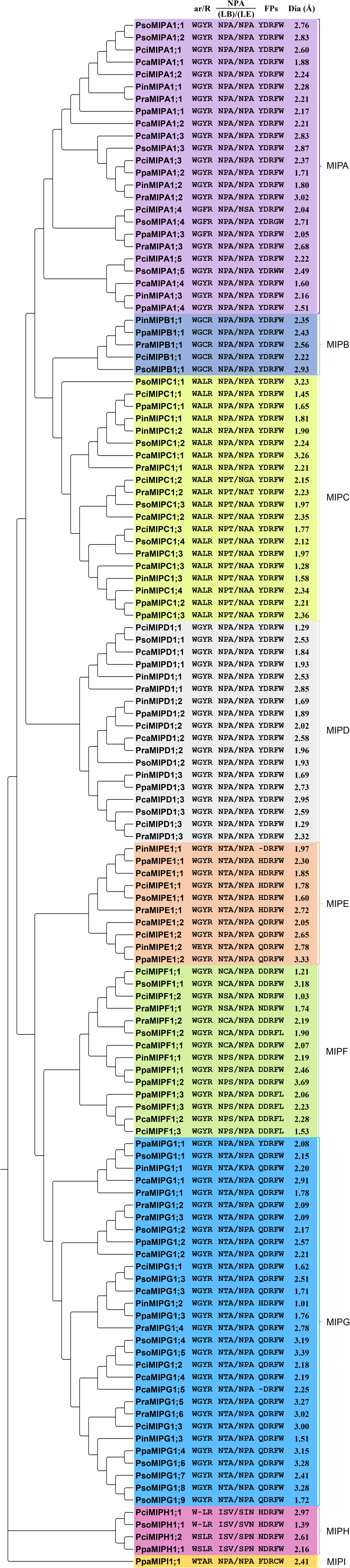
Grouping of MIPs from *Phytophthora* species based on phylogeny showing ar/R selectivity filter, NPA motifs, Froger’s positions, and diameter in the ar/R selectivity filter region. The phylogenetic tree was generated as described in Figure 2. The residues in the ar/R selectivity filter, NPA motifs, and the Froger’s positions were selected from the 3D models, as well as from the alignment shown in Supplementary Figure 3. The pore diameter at the ar/R region, which is considered one of the reasons for transport selectivity, was determined as described in the methods and tabulated on the right side. The different group of PMIPs were indicated with different color and named as MIPA-MIPI, respectively.

### 3.3. Sequence analysis of PMIPs

We calculated the pair-wise sequence identity and similarity among the intragroup and intergroup PMIPs by using EMBOSS Needle (http://www.ebi.ac.uk/Tools/psa/emboss_needle/). The average identity and similarity among the intragroup PMIPs were 76% and 83%, respectively, whereas those in the intergroup PMIPs were 40 and 52%, respectively (Table S2). However, the sequence identity and similarity of MIPHs to MIPA-MIPGs was only 34 and 48%, respectively. Nevertheless, the intergroup average sequence identity among MIPAs-MIPDs and MIPEs-MIPGs was 46 and 50% and similarity was 64 and 62%, respectively. In contrast, the intergroup average sequence identity and similarity of PMIPs from MIPAs-MIPDs to MIPEs-MIPGs was 48 and 63%, respectively. These results indicated that PMIPs of MIPAs-MIPDs were closer compared to those of MIPEs-MIPGs and vice versa. The intergroup sequence identity varied from 2 to 30% and similarity varied from 1 to 56%, indicating that each group was divergent from the others.

### 3.4. Gene structure of Phytophthora MIPs

All the full-length *MIP* sequences found in *P. infestans*, *P. parasitica*, *P. sojae, P. ramorum, P. capsici*, and *P. cinnamomi* were analyzed for introns and exons (Fig. 4). Interestingly, among the 126 PMIPs, 107 homologues had no introns. This characteristic might be common to prokaryotic genes, indicating their origin from prokaryotic homologs. These genes might be introduced to *Phytophthora* genomes by horizontal gene transfer. Thirteen PMIPs had one intron. Two introns were observed in four *PMIP* genes, namely *PciMIPH1;1, PinMIPE1;1, PinMIPG1;2*, and *PsoMIPH1;1*. Three and five introns were found only in *PciMIPA1;4* and *PinMIPC1;3*, respectively. Despite a few disparities, conserved intron positions were not found in *PMIPs* (Fig. 4).

**Fig. 4.**
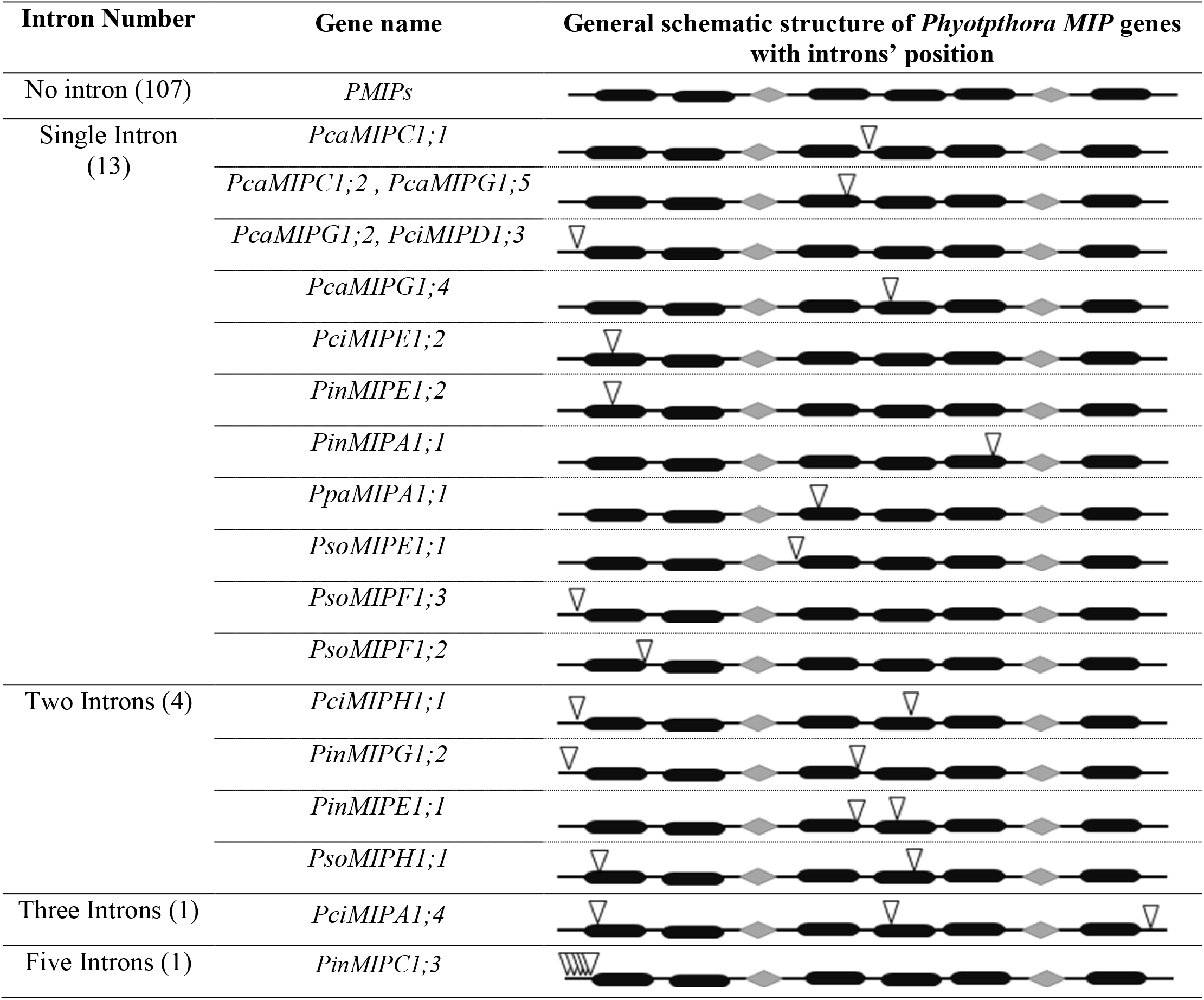
Gene structure of MIPs from *Phytophthora* species. Exon-intron organizations of *MIP* genes from *Phytophthora* species were depicted as described in the methods. The classification of PMIPs based on number of introns mentioned in the first column with the number within the parenthesis indicates the total number of PMIP homologues. *PMIPs* having introns are mentioned in the second column. The six TM regions are shown as black bars and the loops B and E are shown in diamond shapes. The intron positions are indicated by inverted triangles.

Six *PMIP* genes specifically *PcaMIPC1;1*, *PcaMIPC1;1*, *PpaMIPA1;1*, *PpaMIPA1;1*, *PciMIPA1;4* and *PinMIPC1;3* have introns greater than 225 nt including highly repeated sequences. The average length of intron(s) in the remaining 13 PMIPs is 89 nt. which is closer to the average size of fungal introns [65–67]. Short introns are believed to contribute to higher expression of genes in *C. elegans* and *Homo sapiens* [68, 69]. However, the role of intron(s) in *Phytophthora* needs to be determined experimentally.

### 3.5. Analysis of ar/R selectivity filter and Froger’s positions of PMIPs

Amino acid residues involved in the ar/R filter and Froger’s positions are important for functional grouping and substrate selectivity of MIPs [1, 3, 11–17, 70]. The amino acid residues in the ar/R filter and Froger’s positions from the 3D models of all PMIPs were determined. The structural and multiple sequence alignments of PMIPs helped us to identify the residues in the ar/R selectivity filter and the Froger’s positions. The residues at the ar/R selectivity filter and Froger’s positions in nine groups are shown in Fig. 3. Although MIPs in plants and fungi conserve group-specific ar/R selectivity filters and Froger’s positions [1, 3, 8, 11, 15, 20], the ar/R filter and Froger’s positions were identical in several groups of PMIPs (Fig. 3). Despite a few disparities (PciMIPA1;4, PsoMIPA1;4, PpaMIPA1;3, and PinMIPE1;2), the tetrad in the ar/R selectivity filter in MIPAs and MIPDs-MIPGs was WGYR, which was found in ß-AQPGs in fungi [20]. However, MIPBs, MIPCs, MIPHs, and MIPI contained group-specific ar/R selectivity filters, having the tetrad WGCR, WALR, WSLR, and WTAR, respectively, which are not usually observed in MIPs of other taxonomic groups. Nevertheless, the residue in the H5 position of the ar/R filter in PciMIPH1;1 and PsoMIPH1;1 was deleted. The ar/R selectivity filter, as the name suggests, usually consists of an aromatic residue and R, was found in the H2 and LE2 position, respectively. This was true for all PMIPs, although exceptions have been reported in plants, fungi, and algae [1, 8, 15, 20, 21]. The H5 position in PMIPs is occupied by small residues (G/A/S), which is generally the case in many MIPs in plants (TIPs, NIPs, and SIPs), fungi, and algae [1, 8, 20, 21]. However, all PIPs in plants, most of the AQPs in mammals, some groups of MIPs in algae, and bacteria conserved H, and some fungal, algal, archaeal, and plant MIPs conserved L/I/V in the corresponding position [1, 2, 20, 21]. A/C/G/S/T residues are usually found in the LE1 position in MIPs of plants, animals, fungi, algae, archaea, and bacteria [1, 8, 20, 21]. Except for some fungal MIPs, an aromatic residue is not usually available at the LE1 position in eukaryotes. Interestingly, most of the PMIPs (MIPAs, MIPDs-MIPGs) conserved the bulky hydrophobic aromatic Y in the LE1 position (Fig. 3). The bulky aromatic W and Y in the H2 and LE1 positions in many PMIPs in the five groups collectively might have influenced the channel properties and their transport profile. Hydrophobic L, which is not generally found in MIPs of plants, animals, algae, bacteria, and archaea, but available in some fungal MIPs, is conserved in homologues of MIPC and MIPH. This hydrophobic larger amino acid would have changed the channel property and transport profile compared to MIPs that have A/C/G/S/T in the same position. Superposition of the 3D models of PMIPs with crystal structures of bacterial glycerol facilitator (Glps), bovine aquaporin 1 (AQP1), and spinach PIP, SoPIP2;1, or that of intergroup homologues revealed that the architecture of the ar/R selectivity filter in PMIPs would be influenced by the residue at the H5 and LE1 positions (Fig. 5).

**Fig. 5.**
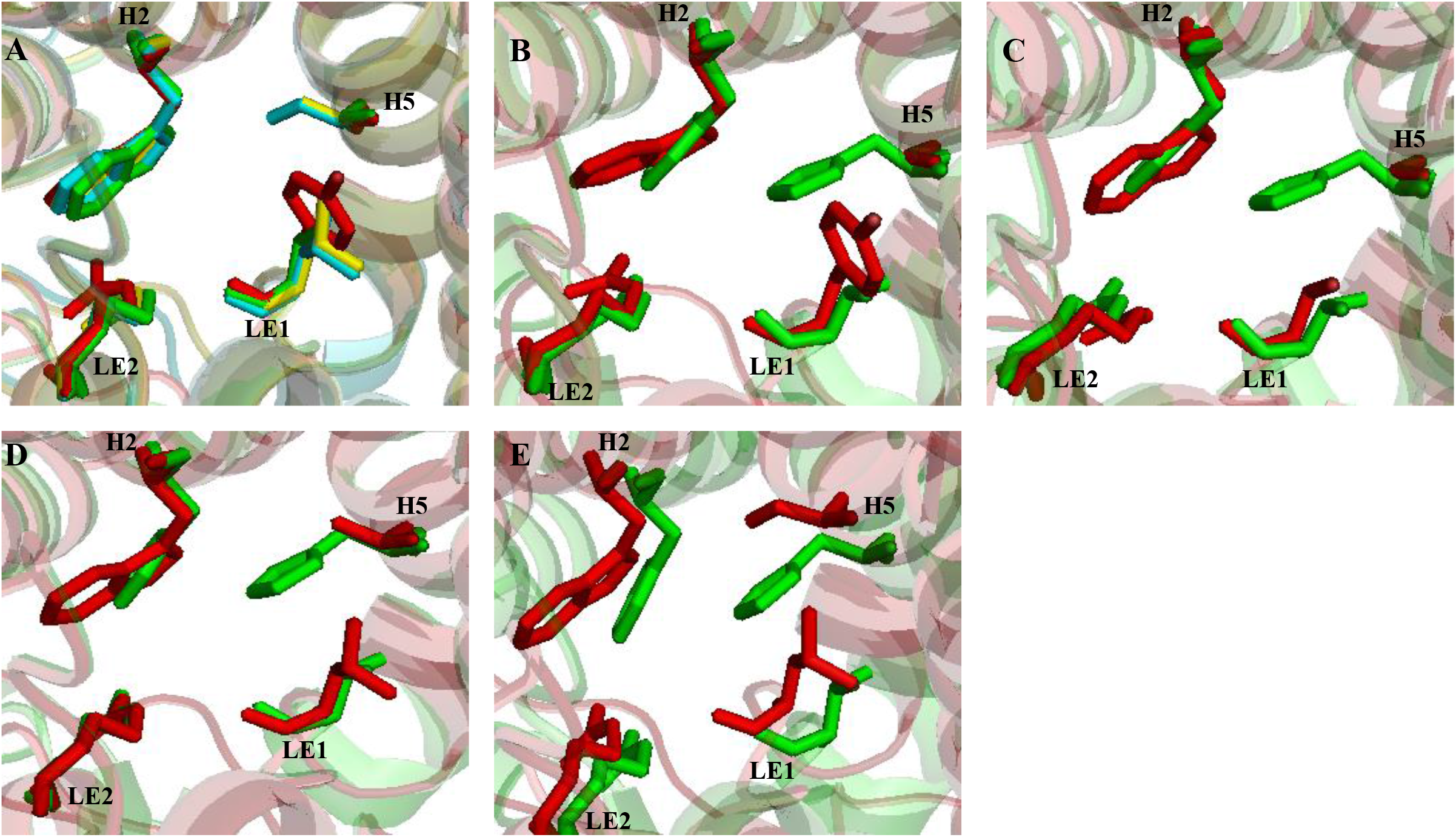
Superposition of 3D models of representative PMIP homologues of group A, B, C, and H. The 3D models of all PMIPs were first constructed separately with SoPIP2;1, and then the models of PciMIPA1;1 (red), PciMIPB1;1 (green), PciMIPC1;1 (yellow), and PciMIPH1;2 (cyan) were superposed (A). The 3D models of PciMIPA1;1 (B), PciMIPB1;1 (C), PciMIPC1;1(D), and PciMIPH1;2 (E) were separately superposed with the template SoPIP2;1. The SoPIP2;1 is shown in green and the PMIP models are in red (B-E). The residues that form the aromatic/arginine (ar/R) tetrad in the superposed structures are shown as sticks. The TM α- helices and the loops to which these residues belong are indicated.

The Froger’s positions, P1 from H3, P2 and P3 from LE, and P4 and P5 from H6 as described by Froger et al. [17] were YDRFW in MIPAs-MIPDs, excluding two homologues, PsoMIPA1;4 and PsoMIPA1;5, in which the F was substituted by G and W, respectively (Fig. 3). Although these Froger’s positions are diverse from MIPs in plants, animals, and algae, most of the groups of fungal MIPs conserve them (Data not shown). In MIPEs-MIPHs, the P2-P5 positions were conserved with DRFW, except PsoMIPF1;2, PpaMIPF1;3, PsoMIPF1;3, PcaMIPF1;2, and PciMIPF1;3, in which the W in the P5 position was substituted by L. The P1 position in these groups was diverse, Q/D in MIPEs; D/N in MIPFs; Q in MIPGs; and H/N in MIPHs. The Froger’s positions in the MIPI (PpaMIPI1;1) was FDRCW. The archaeal and bacterial MIPs reported in Tanghe et al. [55] were analyzed for Froger’s position. The P2-P5 positions were conserved with SALW and TYLW, respectively, and the P1 position was diverse, A/V/L in bacteria and A/V/I in archaea.

### 3.6. PMIPs with substituted NPA motifs

The conserved NPA motifs were found in MIPAs, MIPBs, MIPDs, MIPI, and a subgroup of MIPCs, except PciMIPA1;4, in which the P of the NPA in LE was substituted by S (Fig. 3). As reported for the small basic intrinsic proteins (SIPs) in all plants [1, 4], the homologues of MIPEs-MIPGs conserved a usual NPA motif in the LE, but an unusual NPA motif in the LB, apart from PpaMIPG1;1 that contained the usual NPA in both loops and PinMIPG1;1, in which the N of the NPA in LE was substituted by K. However, unlike SIPs, MIPEs-MIPGs have high molecular weight with longer amino acid sequences. Moreover, although substitution of A by T or L in the NPA motif of LB is observed in SIPs [1, 20], P was substituted by T in all MIPEs and MIPGs. In seven homologues of MIPFs, P was substituted by C or S, and in six homologues of the same group, A was substituted by S. Interestingly, two MIP homologues from each of the six *Phytophthora* species had unusual NPA motifs both in LB and LE, and they were clustered as a subgroup in MIPCs. More intriguingly, the NPA motifs of LB and LE in MIPHs were substituted with ISV and S(P/V/I)N, respectively. Previously, we demonstrated that the NIPs with unusual NPA motifs had characteristic R-rich C-termini [1]. However, MIPCs with both unusual NPA motifs had a RSxGPYE(Y/F) C-terminus followed by the group-conserved GYHH motif. Although MIPHs’ C-termini have no such conserved sequences before the ExQH motif, they contain comparatively more charged residues (Supplementary Fig. 3).

### 3.7. Group-specific consensus sequences/motif of PMIPs

The PMIPs sequences were further compared for group-specific consensus sequences. In the TM regions, the intergroup similarities among the PMIPs were very high. The group-specific deviation was particularly observed in the loops and N- and C-termini (Table 7). Most of the homologues in MIPEs-MIPGs had longer N-termini compared to MIPAs-MIPCs (Data not shown). Interestingly, MIPDs had short N- and C-termini. It is more intriguing that excluding only three PMIPs in group E (PinMIPE1;1, PcaMIPE1;1, and PpaMIPE1;1), all PMIPs conserved a positively charged histidine residue in the C-terminus. With this conserved H residue, all PMIPs conserved a group-specific motif (Table 7). Except MIPEs and MIPHs, the C-termini in all PMIPs were H-enriched, to a greater extent in MIPAs and MIPBs. The C-termini of MIPEs were enriched with D and E and had di- or tri-acidic residues. Despite the NPA motifs in LB and LE, every loop (LA-LE) had group-conserved sequences (Table 7). However, the sequence shown in LB shared both TM2 and LB. In all PMIPs, the LE was longer, where two group-conserved sequences were observed. The first one composed of eight residues included the P2 position. The second consisted of six residues and was located downstream of the P3 position. We further identified group-specific consensus sequences, with one in each of TM3 and TM5, and two in TM4. In the upstream region of TM3, there was an YxxxQ motif in all PMIPs, in which xxx contained group-specific residues. In the consensus sequence composed of eleven residues (Pxxxxxxxxx(M/S) in TM5, the interior positions were blocked with group-wise preserved residues. Nevertheless, the first group-wise conserved sequence located at the start of TM4 was composed of eight residues; and the second one located before the end of the TM4 was composed of seven residues.

## 4. Discussion

In the present study, 126 PMIP homologues were identified from the genomes of six *Phytophthora* species. This study provides comparative information in encoded *PMIP* genes number, diversity of the *PMIP* genes, structural insight of PMIPs, and evolution of PMIPs relative to those in other taxonomic groups.

### 4.1. PMIPs are a new paradigm in microbial aquaporins

Although MIP homologues number varies from organism to organism, plants comparatively possess more homologues than animals and microbes. Although bacterial genomes have 1–2 and fungal and/or algal genomes have only 1–6 *MIP* genes [9, 20, 21, 71], this study showed that the genomes of *Phytophthora* species of oomycetes had 17–27 *MIP* genes (Tables 1–6). The number of *MIP* genes in the genomes of *Phytophthora* species was higher than that even in the human genome and almost similar to that in many plants [1, 5, 18]. The increase in the number of PMIPs might be have been caused by gene duplication event or horizontal gene transfer in addition to the polyploidy nature of *Phytophthora* spp. [72–74]. More interestingly, systematic searching and phylogenetic and structural analysis revealed that PMIPs were distinct and did not cluster to those in other taxonomic groups (Fig. 2, 3). Data collectively indicated that despite some fungal MIPs [20], PMIPs had distinct ar/R filter and Froger’s positions, and substitution in the conserved NPA motifs in comparison with those in plants, animals, algae, bacteria and archaea were divergent. The large numbers, phylogenetical and structural novelty of PMIPs reflected their wide diversity in function and physiological relevance.

### 4.2. Group-specific characteristic C-termini and consensus motifs likely to be associated with novel functions

The uneven length of N- and C-termini of PMIPs groups (Table 7, Supplementary Fig. 3) might have affected their interaction with other molecules or physical interaction for heteromerization of PMIPs [75, 76] because the protein termini were generally exposed on the surface of protein structures making them available for interaction [77, 78]. The heteromerization is one of the mechanisms for regulation of intrinsic permeability of MIPs [76]. The C-termini of proteins are involved in diverse biological processes, such as membrane integration of proteins, protein activity, protein sorting, post-translational modification, protein-protein interaction, or formation of protein complexes [26, 78, 79]. The conserved positively charged H residue in all PMIPs, which is included in the group-specific C-terminal motif (Table 7), is a novel character not observed in other MIPs. Furthermore, despite MIPEs that are enriched with negatively charged D and E residues, the C-termini in most PMIPs are enriched with positively charged H, although its extent is group-specific (Supplementary Fig. 3). Therefore, the distinctive C-termini in PMIPs might be associated with protein sorting, protein-protein interaction, post-translation modification, or other novel functions. KDEL, HDEL, or KKXX motifs in the C-termini of proteins are involved in the retention of protein in the endoplasmic reticulum and stop them to enter into the secretory pathway [78–80]. The NIPs with unusual NPA motifs in LB and LE, have a characteristic R-rich C-termini, which are not seen in NIPs with only one unusual NPA motif, and the R-rich C-termini are thought to be involved in structural stabilization of MIPs [1, 81, 82]. Although a subgroup of MIPCs with both unusual NPA motifs have no such R-rich C-termini, their conserved RSxGPYE(Y/F) sequence would have been involved in the same function in addition to the other aforementioned functions of C-termini because of having charged and phosphorylatable residues. Similarly, MIPHs having both unusual NPA motifs would have different functions because of the presence of comparatively more charged (20–40%) residues in the last 10 amino acids of the C-termini (Supplementary Fig. 3). In contrast, SIPs with only one unusual NPA motif have K-rich C termini [1], which is an important endoplasmic retention signal [4, 81]. However, MIPEs-MIPGs with only one unusual NPA motif have no such K-rich C-termini. It is usually supposed that MIPs with unusual NPA motifs are involved in non-aqua transport rather than water transport [1, 3]. However, it has been demonstrated that two AtSIP1s facilitate transmembrane water diffusion, but not in AtSIP2;1, and it might have non-aqua transport activity [75].

Comparison of PMIPs with MIPs in other taxonomic groups revealed that the group-specific motifs or consensus sequences in PMIPs shown in Table 7 are distinct from the corresponding positions in MIPs of other domains of life (Data not shown). However, some of these motifs or consensus sequences are partly similar to some of the fungal MIPs. Interestingly, the YxxxQ motif in TM3 is largely conserved in most MIPs of other taxonomic groups. This might have important structural roles in MIPs as is reported for NPA motifs. Some of the residues in the group-specific motif or consensus sequences are pore-lining (Supplementary Fig. 3), which in turn, may influence the transport profiles of PMIPs. The substrate-specific signature sequences (SSSS) or specificity-determining positions (SDPs) in the NPA regions, ar/R filters, and Froger’s positions were suggested based on experimentally proven non-aqua facilitator MIPs in plants [1, 8, 70]. These SSSS and SDPs have been shown to be used as tools to predict non-aqua transport activity of plant MIPs [1, 8, 70]. Based on the SSSS or SDPs only in the NPA region of loop B, several PMIPs were predicted to be non-aqua channels (Table S3). However, considering the SSSS or SDPs for three constrictions together, no PMIP was predicted to be non-aqua facilitator unlikely to plant. This result indicated that the SSSS and SDPs of PMIPs in the three constrictions might be structurally distinct, which in turn, would have imparted novel functions to PMIPs. It will be interesting to conduct wet-lab experiments to determine the transport activities and elucidate the structure-function relationship of PMIPs.

## 5. Conclusions

After its discovery in the last decade of the twentieth century, the MIP has fascinated scientists with its potential to aid in understanding the molecular physiology of organisms and development of new innovative pharmacological agents for the treatment of human and plant diseases. It is important to understand host-*Phytophthora* interactions in molecular level. In this context, understanding the insights of evolution, structure, function, and diversity of PMIP channels might be very noteworthy. In the present study, we performed genome-wide identification of MIPs in six *Phytophthora* species, which are devastating plant pathogens and members of oomycetes. Every *Phytophthora* genome encodes several-fold MIP homologues compared to other eukaryotic microbes, such as fungi and algae, even more than in the human genome, and a similar number of homologues as found in many plants. The PMIPs are phylogenetically and structurally distinct from their counterparts in other taxonomic domains. Sequence and 3-D structure analysis studies indicated that the ar/R selectivity filter, group-specific consensus sequences/motifs in different loops and TM helices of PMIPs, and Froger’s positions are distinct from those in other taxonomic domains. The substitutions in the conserved NPA motifs of many PMIPs were also unique. The present study has provided a picture of PMIPs with distinct evolutionary relationships and structural properties. However, we need to perform wet-lab experiments to find out the possible solutes that can pass through the PMIP channels and the physiological relevance of these channel proteins in the *Phytophthora* life cycles.

## Acknowledgements

JA and AH were supported by stipends from The University Grant Commission of Bangladesh. There was no additional external funding received for this study.

## Author’s contributions

AKA and JA conceived the project. AKA designed the work. JA, AH, MAA, MMH and MH carried out the work and participated in analysis with the close supervision of AKA. JA participated in draft preparation of some parts of the manuscript. AKA supervised all procedures and wrote the manuscript. TI and YS critically read the manuscript. All authors approved the final version of the manuscript.

---

**Supplementary Fig. 1.** The phylogenetic tree topology was used to estimate the Ka/Ks ration among branches. Branches marked red showed Ka/Ks ratio >1. Branches were drawn in proportion to their lengths, estimated using the one-ratio model (M0).

**Supplementary Fig. 2. Grouping of MIPs from *Phytophthora* species based on phylogeny.** The phylogenetic tree was generated as described in Figure 2. Each MIP group is shown with a specific background color to distinguish them from others.

**Supplementary Fig. 3. Multiple sequence alignment of MIPs in the six *Phytophthora* species.** The amino acid sequences were aligned using the Clustal Omega program. Each MIP group is shown with a specific background color to distinguish them from others. The TM helices are shown within boxes with black lines and the dual NPA motifs are shown as gray. The residues in the ar/R selectivity filter and FP are red and blue boxed, respectively. The region of the TM helices and loops from which consensus sequences or motifs were depicted in Table 7 are green boxed. The pore-lining residues are indicated by arrows above the alignment and the conserved residues are indicated by stars (*) at the bottom of the alignment.

**Table S1:** Predicted sub-cellular localization of PMIPs.

**Table S2:** Average pairwise sequence identity and similarity between different PMIP groups.

**Table S3:** Prediction of non-aqua substrate transport activity of PMIPs. Based on the signatures sequences in the regions of ar/R filter, Froger’s positions and loops B and E showed for plant aquaporins [1, 8] have been conducted on PMIPs. Considering the signature sequences at ar/R filter, Forger’s positions and loop E, no PMIP had been predicted to facilitate non-aqua substrate as listed. Only based on the signature sequence on loop B, some PMIPs that were predicted to facilitate non-aqua substrates had been listed.

